# Fluidity of gender identity induced by illusory body-sex change

**DOI:** 10.1101/2020.01.13.905315

**Authors:** P. Tacikowski, J. Fust, H. H. Ehrsson

## Abstract

Gender identity is the inner sense of being male, female, both, or neither. How this sense is linked to the perception of one’s own masculine or feminine body remains unclear. Here, in a series of three behavioral experiments conducted on a large group of healthy volunteers (N=140), we show that a perceptual illusion of having the opposite-sex body was associated with a shift toward more balanced identification with both genders and less gender-stereotypical beliefs about one’s own personality characteristics, as indicated by subjective reports and implicit behavioral measures. These findings demonstrate that the ongoing perception of one’s own body affects the sense of one’s own gender in a dynamic, robust, and automatic manner.

## Introduction

Gender identity is the inner sense of being male, female, both, or neither^1,2^. This sense has a profound impact on our life, but little is known about how gender identity is formed and maintained. Apart from its general relevance to all of us as individuals with a sense of gender, an understanding of the neurocognitive mechanisms of gender identity is important for the treatment of gender dysphoria, which is characterized by prolonged distress due to inconsistency between an individual’s biological sex and subjectively perceived own gender^2–4^.

In the past, gender identity was conceptualized as a male-female dichotomy, but current theories often postulate that the sense of own gender is a spectrum of associations with the male and female genders^1,2,5–7^. There is also a general consensus in the field that gender identity is a multifaceted phenomenon that consists of biological, affective, cognitive, and sociocultural components^1,2,5–7^; what exactly these components are and what role they play are matters of debate. Clinical observations suggest that the perception of one’s own body is one of the key aspects of the sense of own gender. For example, people with gender dysphoria (see earlier) often avoid looking in the mirror, hide their body under loose-fitting clothes, and seek hormonal and/or surgical treatment to adjust their physical appearance to their subjective sense of gender^2–4^. Moreover, among cisgender individuals, whose gender identity matches their biological sex, mastectomy and androgen deprivation therapy, which both involve changes of feminine or masculine bodily characteristics, are often related to a crisis of gender identity^8,9^. There are also data suggesting that the mental image of one’s own body is diffused in transgender individuals^10^ and that the brain regions involved in various aspects of body representation are anatomically and functionally atypical in this group^11–14^. However, the link between own body perception and gender identity remains poorly understood from a behavioral experimental perspective, and we do not know whether, and if so how, the perceived sex of one’s own body influences the sense of gender identity in nontransgender individuals.

The full-body ownership illusion^15^ is a powerful experimental tool to manipulate the perception of one’s own body^16–21^. During this illusion, the participants wear head-mounted displays (HMDs) and observe a stranger’s body from the natural first-person perspective. The stranger’s body is continuously stroked with a stick or a brush, and the experimenter applies synchronous touches on the corresponding parts of the participant’s body, which is out of view. This synchronous visuotactile stimulation induces a feeling that the stranger’s body is one’s own, whereas asynchronous stimulation breaks the illusion and serves as a well-matched control condition^15,22–24^. The full-body ownership illusion, as well as the similar rubber hand illusion that involves a single limb^25–28^, occurs when visual, tactile, proprioceptive, and other sensory signals from the body are combined at the central level into a coherent multisensory representation of one’s own body^15–17,21^. Body ownership illusions involving limbs^28^ and full bodies^22,24,29^ are related to increased neural activity in regions of the frontal and parietal association cortices that are related to multisensory integration, such as the ventral premotor and intraparietal cortices. Because these brain regions contain trimodal neurons that integrate visual, tactile, and proprioceptive signals, and because body illusions closely follow the temporal and spatial constraints of multisensory integration, it was proposed that combining bodily signals from different modalities is a key mechanism of attributing ownership to one’s own body^16–21^. The full-body ownership illusion has been replicated in numerous studies^15,22–24,30–36^, even with bodies of the opposite sex^15,35^, but the cognitive consequences of this transient physical sex change on the sense of own gender have not been assessed.

Here, we conducted three within-subject behavioral experiments on a total of one hundred forty naïve healthy volunteers to investigate a possible dynamic relationship between the perception of one’s own body and various aspects of gender identity. In all three experiments, we induced the perceptual “body-sex-change illusion,” which is analogous to the standard full-body illusion (see earlier), but in the HMDs, the participants saw the opposite-sex stranger’s body (Movies S1 and S2). We hypothesized that if one’s own body perception dynamically shapes gender identity, then even a brief transformation of one’s own perceived physical sex during the body-sex-change illusion should shift the affective, implicit, and personality-related aspects of gender identification toward those of the opposite gender.

## Results

Experiment I tested whether the perception of one’s own body dynamically shapes the subjective feeling of masculinity/femininity. The experiment comprised a 2-by-2 factorial design with four conditions (Fig. 1a): “synchronous opposite sex” (syncO), “synchronous same sex” (syncS), “asynchronous opposite sex” (asyncO), and “asynchronous same sex” (asyncS). This design allowed us to manipulate the sex-related characteristics of one’s perceived bodily self in the body-sex-change illusion condition (syncO) while controlling for potential confounding factors related to experiencing the body ownership illusion per se (syncS) or cognitive biases due to simply looking at a male or female body (asyncS, asyncO). We measured the illusion psychometrically, by asking the participants to rate their subjective experience of owning the stranger’s body (Fig. 1b), and objectively, by recording the participants’ physiological fear reactions (skin conductance responses) when the stranger’s body was physically threatened with a knife (Fig. 1c). Both of these illusion measures should be higher during the synchronous than during the asynchronous conditions^15,23,29,31,34^. Importantly, before experiencing any body perception manipulation (baseline) and after every full-body illusion condition, the participants rated how masculine or feminine they felt (Fig. 1d and 1e).

**Fig. 1.**
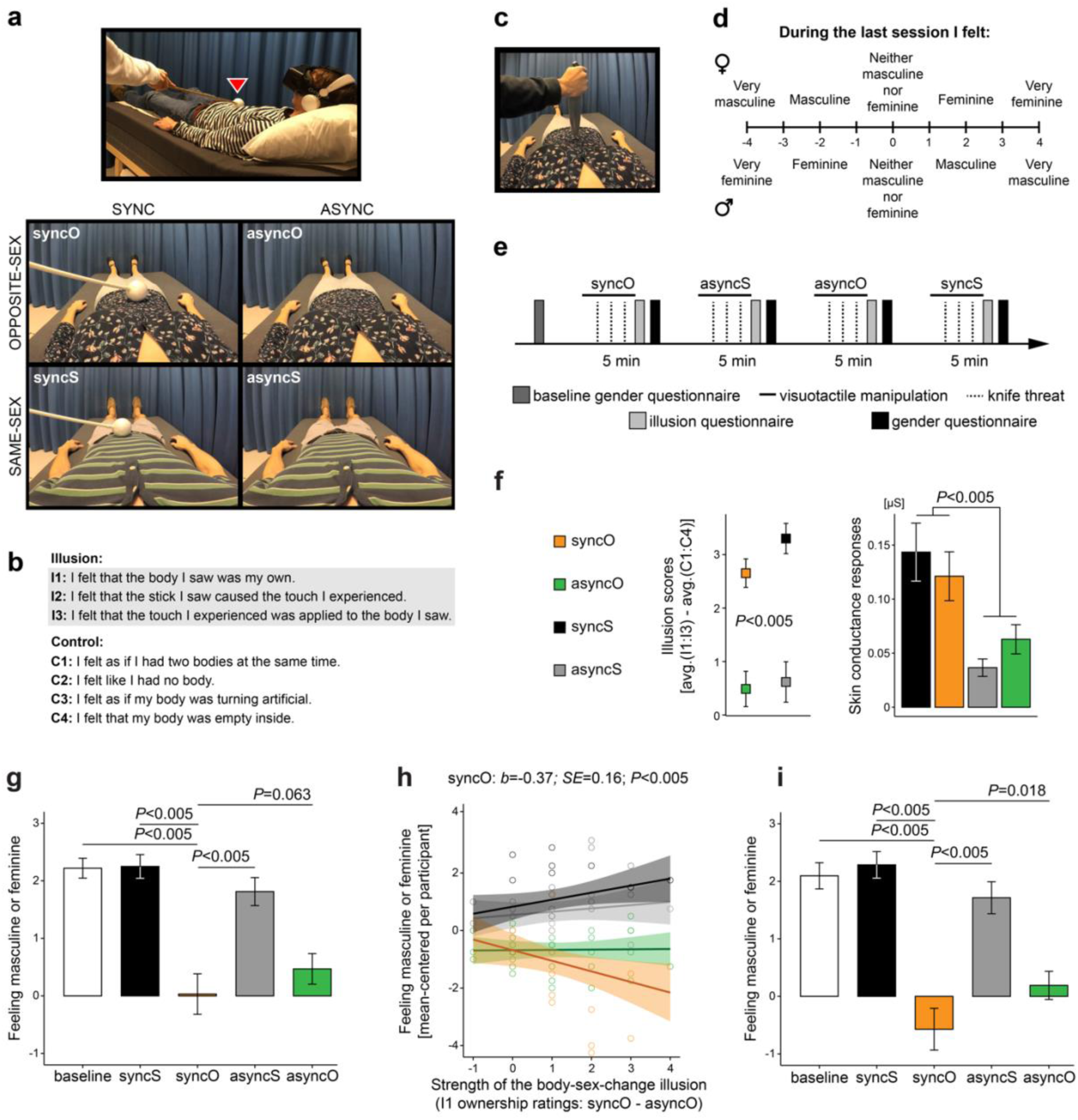
Perceptual illusion of having the opposite-sex body modulated the subjective experience of feeling masculine or feminine (Experiment I). **(a)** The participants (N=32; 15 females) lay on a bed and wore a head-mounted display in which a body of an unknown male or female was shown from a first-person perspective (the participant’s real body was out of view). Video frames illustrate all four conditions for a male participant (top picture). For a female participant, the videos from the lower and upper rows would be swapped. In the synchronous conditions, touches applied to the participant and touches applied to the stranger’s body were matched (see the red triangle), whereas in the asynchronous conditions, touches applied to the participants were delayed by 1 s. We expected to induce the body-sex-change illusion specifically in the syncO condition, and the other conditions served as controls. **(b)** After each condition, the participants rated illusion (I1:I3) and control (C1:C4) statements on a 7-point scale (−3 – “strongly disagree”; +3 – “strongly agree”). The illusion statements assessed the feeling that the stranger’s body is one’s own, whereas the control statements controlled for any potential effects of suggestibility or task compliance. **(c)** Genuine ownership of the stranger’s body should be associated with increased physiological stress responses of the participant when the stranger’s body is physically threatened. Thus, we measured the participants’ skin conductance responses elicited by brief “knife threat” events that occurred in the videos. **(d)** Before the experiment (baseline) and after each condition, the participants rated how feminine or masculine they felt. The upper row shows scale assignment for female participants and the lower row for males. **(e)** The order of the conditions was counterbalanced across the participants, and the whole experiment lasted ∼30 min. **(f)** The illusion ratings and the magnitude of skin conductance responses were significantly higher in the synchronous than in the asynchronous conditions, which shows that the full-body ownership illusion was elicited as expected. **(g)** During syncO, the female participants indicated feeling less feminine, and the male participants indicated feeling less masculine than during other conditions. **(h)** Strong illusory ownership of the opposite-sex body in syncO was related to a significant shift toward the opposite gender. **(i)** The participants who experienced a strong body-sex-change illusion (above-median I1 ownership ratings: syncO – asyncO; N=21) indicated feeling more like the opposite gender during syncO. Plots show means ± *SE*.

The full-body ownership illusion was induced as expected; that is, the participants agreed more with illusion statements in the synchronous than in the asynchronous conditions, and knife threats that occurred during the synchronous conditions triggered stronger skin conductance responses than knife threats that occurred during the asynchronous conditions (Fig. 1f; main effect of synchrony; illusion questionnaire: *b*=2.68; *SE*=0.31; *t*=8.59; *P*<0.005; 95% CI [2.1, 3.3]; skin conductance: *b*=0.1; *SE*=0.02; *t*=5.07; *P*<0.005; 95% CI [0.06, 0.14]; linear mixed models; two-sided; N=32). With regard to our main hypothesis, we found that during the syncO condition, the female participants indicated feeling less feminine and the male participants indicated feeling less masculine than during the baseline, syncS, and asyncS control conditions (Fig. 1g; synchrony × body interaction; *b*=−0.88; *SE*=0.4; *t*=−2.16; *P*=0.028; 95% CI [−1.66, −0.1]; pairwise comparisons: syncO vs. baseline: *b*=−2.19; *SE*=0.35; *t*=−6.22; *P*<0.005; 95% CI [−2.88, −1.52]; syncO vs. syncS: *b*=−2.22; *SE*=0.37; *t*=−6.06; *P*<0.005; 95% CI [−2.94, −1.52]; syncO vs. asyncS: *b*=−1.78; *SE*=0.37; *t*=−4.83; *P*<0.005; 95% CI [−2.48, −1.05], linear mixed models; two-sided; N=32). The difference between syncO and asyncO was a significant trend in the hypothesized direction (*b*=−0.44; *SE*=0.23; *t*=−1.87; *P*=0.063; 95% CI [−0.88, 0.02]; linear mixed model; two-sided; N=32). Importantly, the shift toward the opposite gender during syncO was enhanced by the illusory ownership of the opposite-sex body, operationalized as the difference between I1 ownership ratings in syncO – asyncO (Fig. 1h; synchrony × body × ownership interaction; *b*=−0.51; *SE*=0.24; *t*=−2.1; *P*=0.037; 95% CI [−0.98, −0.04]; linear mixed model; two-sided; N=32; follow-up analyses of the main effect of ownership in each condition; syncO: *b*=−0.37; *SE*=0.16; *t*=−2.38; *P*<0.005; 95% CI [−0.69, −0.13]; syncS: *b*=0.24; *SE*=0.12; *t*=1.96; *P*=0.028; 95% CI [0.03, 0.46]; asyncO: *b*=0.01; *SE*=0.09; *t*=0.11; *P*=0.9; 95% CI [−0.17, 0.2]; asyncS: *b*=0.11; *SE*=0.12; *t*=0.94; *P*=0.26; 95% CI [−0.08, 0.35]; baseline: *b*=0.04; *SE*=0.15; *t*=0.32; *P*=0.68; 95% CI [−0.19, 0.3]; linear regression models; two-sided; N=32). Moreover, the participants who experienced a strong body-sex-change illusion (N=21; median-split; see Methods) indicated feeling more like the opposite gender in syncO than in the other conditions (Fig. 1i; synchrony × body interaction; *b*=−1.33; *SE*=0.54; *t*=−2.49; *P*=0.014; 95% CI [−2.37, −0.27]; pairwise comparisons: syncO vs. baseline: *b*=−2.67; *SE*=0.42; *t*=−6.38; *P*<0.005; 95% CI [−3.48, −1.84]; syncO vs. syncS: *b*=−2.86; *SE*=0.42; *t*=−6.82; *P*<0.005; 95% CI [−3.67, −2.02]; syncO vs. asyncS: *b*=−2.29; *SE*=0.45; *t*=−5.13; *P*<0.005; 95% CI [−3.17, −1.42]; syncO vs. asyncO: *b*=−0.76; *SE*=0.33; *t*=−2.31; *P*=0.018; 95% CI [−1.41, −0.12]; linear mixed models; two-sided; N=21). Overall, Experiment I shows that the ongoing perception of one’s own body dynamically updates the subjective feeling of masculinity or femininity.

Experiment II tested whether the perceived sex of one’s own body also modulates the *implicit* structure of associations between oneself and both genders. This experiment had the same two-by-two factorial design as Experiment I (Fig. 1a and 2a), but this time, gender identity was measured with the Implicit Association Test (IAT)^37–41^. During this test, the participants heard words belonging to four semantic categories (*male*, *female*, *self*, *other*) and sorted these words into just two response categories. In one block, the participants responded with the same key to words from the *self* and *female* categories, which made this block congruent for females and incongruent for males. In the other block, the participants responded with the same key to words from the *self* and *male* categories, which made this block incongruent for females and congruent for males (Fig. 2b). The congruent block is typically related to shorter reaction times (easier task) than the incongruent block (harder task), and the difference in the responses from the two blocks provides an objective behavioral proxy of a person’s gender identification^37–41^. The participants performed the IAT four times, once *during* each condition, which allowed us to track changes in the implicit structure of gender identification across different embodiment contexts (Fig. 2c).

**Fig. 2.**
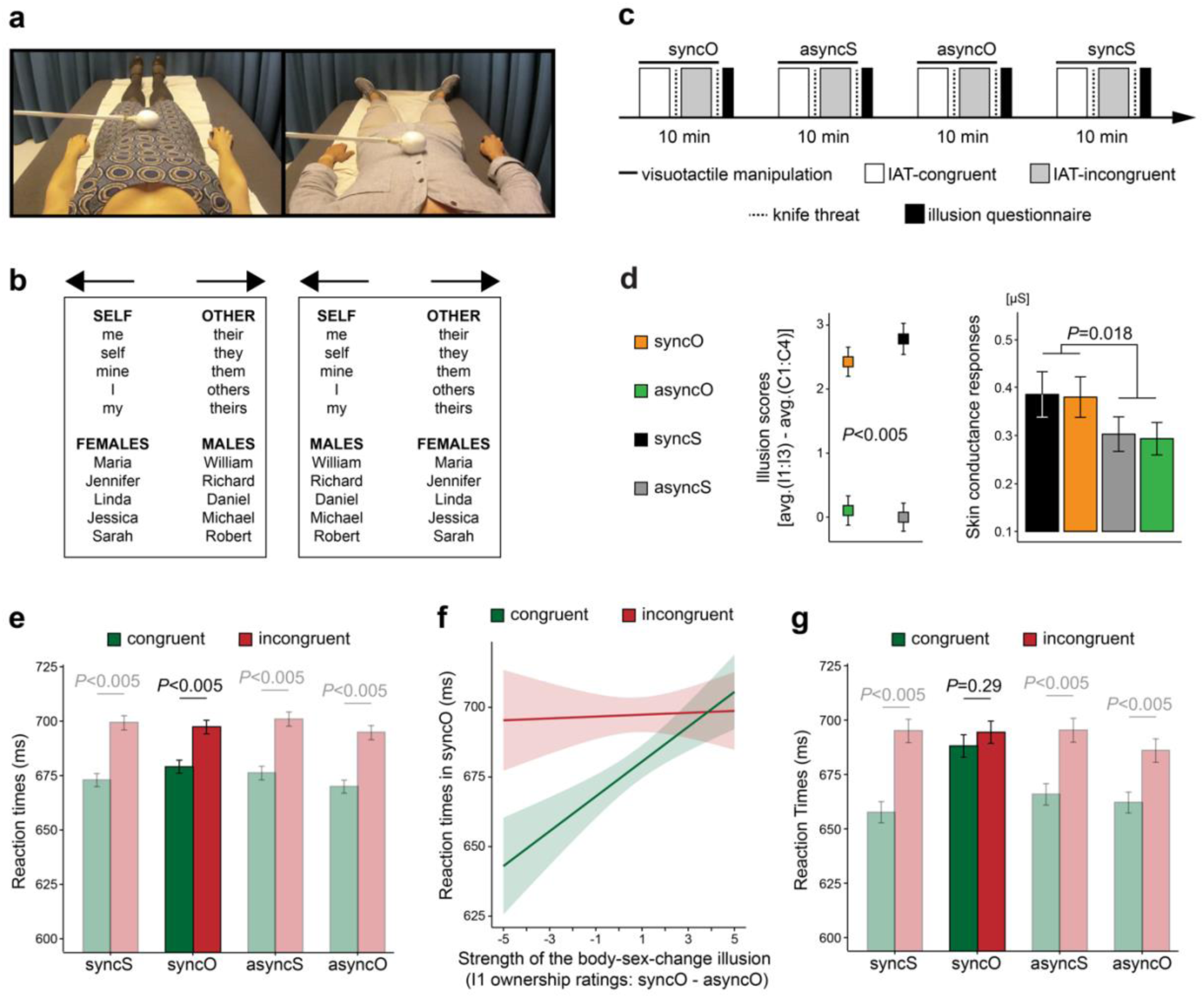
The body-sex-change illusion balanced implicit associations between the self and both genders (Experiment II). **(a)** This experiment (N=64; 32 females) comprised the same four conditions as Experiment I (Fig. 1a), but we used recordings of a different male and female body to enhance the generalizability of our findings. **(b)** In one IAT block (left panel), the participants responded with the same key (indicated by the arrows) to words from the *self* and *female* categories, which made this block congruent for females and incongruent for males. In the other block (right panel), the participants responded with the same key to words from the *self* and *male* categories, which made this block incongruent for females and congruent for males. **(c)** During each condition, the participants completed one full IAT (two blocks). The condition order and block order were counterbalanced across participants, and the whole experiment lasted ∼ 60 min. **(d)** The body-sex-change illusion was successfully induced, as shown by questionnaire data and the magnitude of threat-evoked skin conductance responses. **(e)** In all conditions, reaction times were significantly shorter in the congruent than in the incongruent blocks, which shows that it was generally easier for the participants to associate themselves with the gender category consistent with their sex. **(f)** Strong illusory ownership of the opposite-sex body in syncO was related to the balancing of implicit associations between the self and both genders, and this effect was related mainly to a weakening of the associations between the self and preferred gender (i.e., a lengthening of the reaction times in the congruent block). For clarity of display, individual data points are not shown (n=7290). **(g)** The participants who experienced a strong body-sex-change illusion (above-median I1 ownership ratings: syncO – asyncO; N=24) responded similarly quickly in the incongruent and congruent blocks during syncO. Bar plots show means ± *SE*.

The body-sex-change illusion was also successfully induced in Experiment II, as demonstrated by both the questionnaire and skin conductance data (Fig. 2d; main effect of synchrony; illusion questionnaire: *b*=2.78; *SE*=0.24; *t*=11.51; *P*<0.005; 95% CI [2.31, 3.26]; skin conductance: *b*=0.08; *SE*=0.04; *t*=2.38; *P*=0.018; 95% CI [0.014, 0.153]; linear mixed models; two-sided; N=64). In all conditions, the participants found it easier to associate themselves with the gender that was consistent with their sex, as indicated by shorter reaction times in the congruent than in the incongruent blocks (Fig. 2e; main effect of block; *b*=25; *SE*=4; *t*=6.53; *P*<0.005; 95% CI [18, 33]; follow-up analyses of the main effect of block in each condition: syncO: *b*=18; *SE*=4; *t*=4.85; *P*<0.005; 95% CI [11, 26]; syncS: *b*=28; *SE*=4; *t*=7.35; *P*<0.005; 95% CI [21, 36]; asyncO: *b*=25; *SE*=4; *t*=6.68; *P*<0.005; 95% CI [18, 32]; asyncS: *b*=25; *SE*=4; *t*=6.51; *P*<0.005; 95% CI [18, 33]; linear mixed models; two-sided; N=64). Importantly, the strong illusory ownership of the opposite-sex body in syncO was related to a reduced difference between the incongruent and congruent blocks, which shows that the illusion balanced the strength of implicit associations between the self and both genders (Fig. 2f; synchrony × body × block × ownership interaction; *b*=14; *SE*=4; *t*=3.76; *P*<0.005; 95% CI [7, 21]; follow-up analysis of block × ownership interaction in syncO: *b*=−6; *SE*=2; *t*=−3.23; *P*<0.005; 95% CI [−9, −2]; linear mixed models; two-sided; N=64). The control analyses showed that there was no significant reduction between the reaction times in the congruent vs. the incongruent blocks in other conditions (block × ownership interaction; syncS: *b*=6; *SE*=2; *t*=3.38; *P*=0.99; 95% CI [3, 10]; asyncS: *b*=0; *SE*=2; *t*=0.04; *P*=0.52; 95% CI [−4, 4]; asyncO: *b*=2; *SE*=2; *t*=1.08; *P=*0.86; 95% CI [−2, 5]; linear mixed models; one-sided; N=64). Interestingly, this balancing of implicit associations with both genders in syncO was related to a lengthening of the reaction times in the congruent block rather than to a shortening of the reaction times in the incongruent block, which suggests that the body-sex-change illusion mainly hindered the existing associations between the self and gender category consistent with one’s sex (Fig. 2f; main effect of ownership; syncO-congruent: *b*=6; *SE*=1; *t*=4.33; *P*<0.005; 95% CI [3, 9]; syncO-incongruent: *b*=0; *SE*=2; *t*=0.25; *P*=0.81; 95% CI [−3, 3]; linear regression models; two-sided; N=64). Furthermore, the participants who experienced strong illusory ownership of the opposite-sex body in syncO (N=24; median-split; see Methods) had similar reaction times in the congruent and incongruent IAT blocks (Fig. 2g; sync × body × block interaction; *b*=27; *SE*=13; *t*=1.99; *P*=0.046; 95% CI [0.4, 53]; follow-up analyses of the main effect of block in each condition; syncO: *b*=7; *SE*=7; *t*=1.03; *P*=0.29; 95% CI [−6, 19]; syncS: *b*=39; *SE*=7; *t*=5.9; *P*<0.005; 95% CI [26, 52]; asyncO: *b*=25; *SE*=7; *t*=3.88; *P*<0.005; 95% CI [13, 38]; asyncS: *b*=31; *SE*=7; *t*=4.62; *P*<0.005; 95% CI [17, 44]; linear mixed models; two-sided; N=24). Thus, the main finding of Experiment II is that the moment-to-moment perception of one’s own body balanced the strength of the implicit associations between the self and both gender categories.

Apart from the basic categorization (“I am a man/woman”), gender identity includes stereotypical gender roles, for example, “I am emotional” for females, or “I am competitive” for males^2,7,37^. These roles constitute what it means for a given person to be male or female in a given sociocultural context and are important in our lives, but they can also prevent some individuals from realizing their full personal or professional potential. Experiment III investigated whether the body-sex-change-induced fluidity of gender identity is generalized to stereotypical gender roles. This experiment consisted of two conditions: syncO and asyncO; thus, in the HMDs, the female participants always observed a male body, and the male participants always observed a female body (Fig. 3a and 3b). After each condition, the participants filled out a short version of the Bem Sex-Role Inventory (BSRI)^42,43^ that contained personality characteristics that are stereotypically associated with the male and female genders (Fig. 3b and 3c). The participants’ task was to rate how well they thought each characteristic described them.

**Fig. 3.**
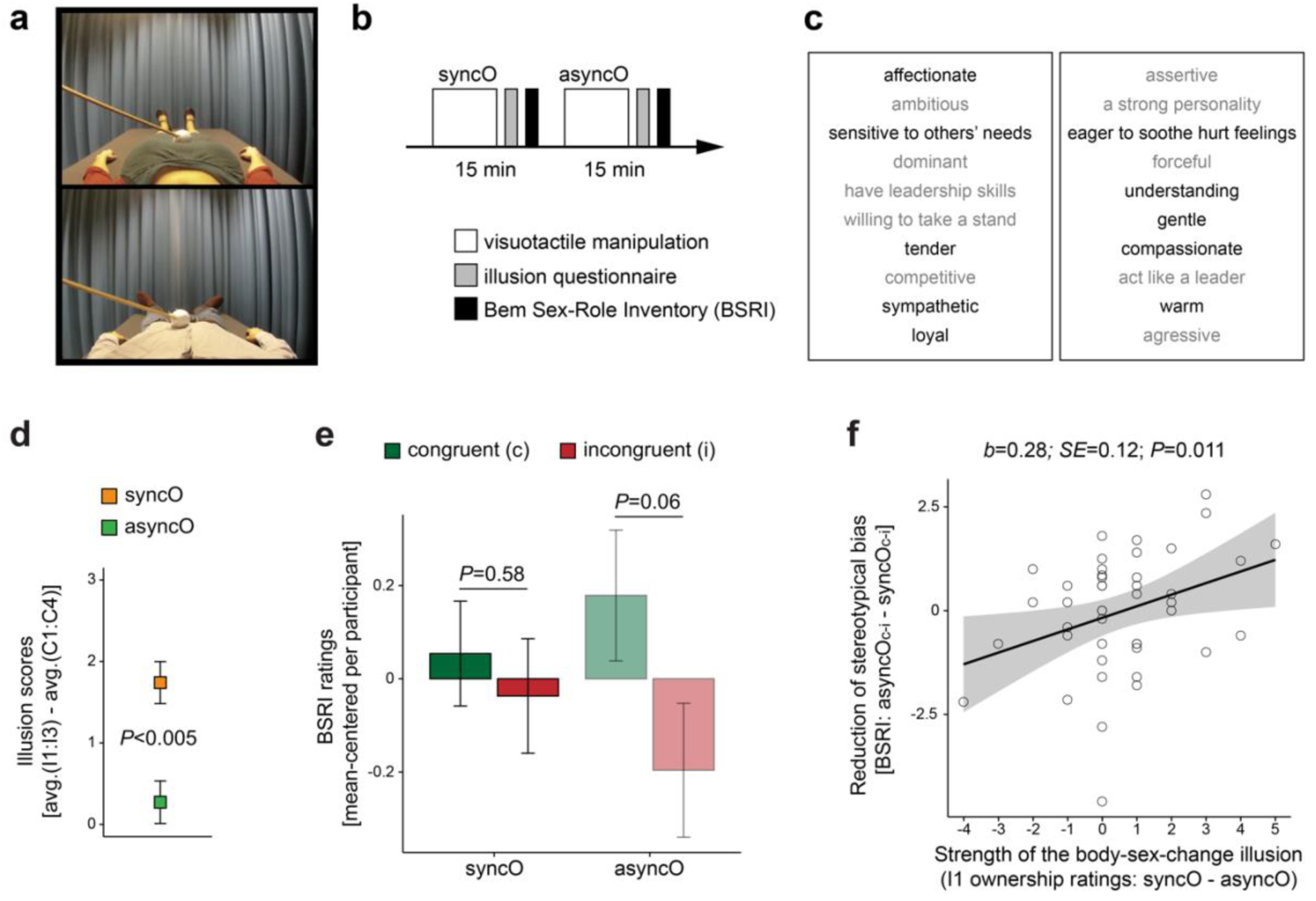
Illusory ownership of the opposite-sex body was associated with less gender-stereotypical beliefs about one’s own personality traits (Experiment III). **(a)** Frames from the videos used in this experiment (N=44; 22 females). **(b)** The experiment consisted of two conditions (syncO and asyncO); thus, in the head-mounted display, the female participants always observed a male body and the male participants a female body that was stroked either synchronously or asynchronously with regard to touches delivered to the participants. The condition order was counterbalanced across the participants, and the whole experiment took ∼ 45 min. **(c)** After each condition, the participants rated how well each personality characteristic describes the self (1 – “not at all”; 7 – “very much”). Each BSRI sublist contained five traits stereotypically related to males (gray) and five traits stereotypically related to females (black). **(d)** The illusion ratings were significantly higher in the syncO condition than in the asyncO condition, which demonstrates that the body-sex-change illusion was efficiently induced. **(e)** In the asyncO (control) condition, the participants tended to indicate gender-stereotypical beliefs about their own personality, whereas in the syncO body-sex-change illusion condition, this bias was reduced. **(f)** Importantly, strong illusory ownership of the opposite-sex body was related to less stereotypically biased gender roles. Bar plots show means ± *SE*.

We found that the body-sex-change illusion was efficiently induced in Experiment III as well (Fig. 3d; main effect of synchrony; illusion questionnaire: *b*=1.47; *SE*=0.25; *t*=5.99; *P*<0.005; 95% CI [0.98, 1.96]; linear mixed model; two-sided; N=44). In the asyncO control condition, the participants tended to indicate stereotype-congruent beliefs about their own personality, whereas in the syncO condition, this bias was reduced, as indicated by similarly high self-ratings of stereotypically masculine and feminine traits (Fig. 3e; stereotype-congruent vs. stereotype-incongruent; asyncO: *b*=0.37; *SE*=0.2; *t*=1.89; *P*=0.062; 95% CI [−0.02, 0.77]; syncO: *b*=0.09; *SE*=0.16; *t*=0.55; *P*=0.58; 95% CI [−0.42, 0.24]; linear mixed models; two-sided; N=44). Importantly, strong illusory ownership of the opposite-sex body was related to less biased beliefs about one’s own gender roles (Fig. 3f; main effect of ownership; *b*=0.28; *SE*=0.12; *t*=2.38; *P*=0.011; 95% CI [0.07, 0.47]; linear regression model; two-sided; N=44). These findings show that the perception of one’s own masculine or feminine physical characteristics flexibly updates stereotypical beliefs about one’s own personality.

Finally, to assess the overall robustness of the relationship between own body perception and gender identity, we performed a post hoc meta-analysis on data from all three experiments combined. We found that strong illusory ownership of the opposite-sex body in syncO was related to increased updating of the sense of own gender, as measured in the experiments (Supplementary Fig. S1; *ρ*^138^=0.24; *P*<0.005; Spearman correlation; two-tailed). Control analyses on the data from all three experiments further showed that the male and female participants experienced illusory ownership of the opposite-sex body equally strongly and that there was no consistent significant relationship between the body-sex-change illusion strength and the participants’ age or baseline masculinity/femininity ratings (Supplementary Fig. S2; for similar evidence during syncS, see Supplementary Fig. S3). Moreover, the degree of gender identity updating did not significantly differ between males and females and was not significantly related to the participants’ age or baseline masculinity/femininity ratings (Supplementary Fig. S4). These results are in line with the notion that the full-body illusion is a robust multisensory perceptual phenomenon that generally is not affected by high-level cognitive or emotional processes^16,21^, which validates the current illusion-based approach to dynamically changing the perceptual basis of the bodily self in a mixed group of male and female subjects.

## Discussion

The present study used the body-sex-change illusion to experimentally investigate the link between own body perception and gender identity. We found that even a brief transformation of one’s own perceived bodily sex dynamically updated the affective, implicit, and personality-related aspects of the sense of own gender and made these aspects more balanced across masculine and feminine categories. This main finding was consistent across three separate experiments conducted on a large group of healthy volunteers with the use of subjective and objective behavioral measures. The remarkable fluidity of gender identity that we report here extends previous knowledge by demonstrating that the link between own body perception and the sense of own gender is dynamic, robust, and direct. It is dynamic because the effects that we detected occurred within minutes of the body-sex-change illusion duration, it is robust because these effects were present at explicit and implicit levels, and it is direct because the changes in gender identity precisely followed our experimental manipulation of perceived own body sex.

The above conclusions have important implications for existing theories of gender identity development. Many of these theories focus entirely on cognitive and social factors and neglect the importance of own body perception (for reviews, see^44,45^). In contrast, our findings provide clear evidence that the representation of one’s own body is an integral part of the sense of own gender. Thus, apart from the conceptual components, gender identity seems to consist of representations that are sensorimotor and nonverbal in nature; these embodied representations are presumably acquired largely unconsciously, from early infancy onwards, by extracting patterns of one’s own and other people’s touch, movement, and affective reactions^5,6^. Another common theoretical assumption that our results challenge is that the development of gender identity requires gender constancy, that is, a child’s realization that (i) they are a boy or a girl, (ii) this identity does not change over time, and (iii) this identity is not affected by gender-typed appearance or activities (for reviews, see^44,45^). Instead, we demonstrate that the affective, implicit, and personality-related aspects of gender identity are continuously updated by the moment-to-moment representation of one’s own body. This finding does not deny the obvious fact that most people have a stable gender identity but suggests that this stability depends on coherence among input variables in the gender identity construction process. The above claim is based on the idea that the sense of own gender emerges from dynamic coupling between relevant biological, psychological, and sociocultural factors; when all these factors cohere tightly, gender identity remains stable, but when coherence is poor, gender identity is updated accordingly^5,6^. The present findings demonstrate that a change in the appearance of one’s own body can sufficiently perturb the sense of own gender.

A thorough reader might ask: if the perception of one’s own body is so critical for gender identity, then how can these two aspects remain in conflict for a prolonged period of time in transgender individuals? First, our results should not be treated as evidence that the perceived bodily sex is the *only* factor that shapes the sense of own gender; instead, this sense is a complex phenomenon that is constructed from multiple factors^1,2,5–7^. Second, some symptoms of gender dysphoria, such as avoiding looking in the mirror or hiding one’s body under loose-fitting clothes^2–4^, actually suggest that these patients might actively suppress the link between their own body perception and their subjective sense of gender. Our results contribute to the discussion about gender identity development by suggesting that there is a continuous bottom-up influence on cognitive, conceptual, and affective representations of gender identity from the perceptual body representation in terms of secondary sexual characteristics. Future studies should investigate the important question of how transgender individuals, with and without gender dysphoria, update their sense of own gender during the body-sex-change illusion.

Another key finding of the present study is that the body-sex-change illusion reduced gender-stereotypical beliefs about one’s own personality. This result suggests that gender identification (“I’m male/female”), gender stereotypes (e.g., “Females are emotional”, “Males are competitive”), and stereotypical gender roles (e.g., “I’m emotional/competitive”) are connected with each other^2,37^ so that a change in one aspect (gender identification), due to the body-sex-change illusion, affects other aspects (stereotypical gender roles). It is worth mentioning that existing programs against gender discrimination, such as media campaigns or educational workshops, target mainly explicit manifestations of gender stereotypes^46^. However, people are often unaware of the fact that their way of thinking is biased, and thus they cannot deliberately change it. Body-oriented techniques, similar to the one used here, could possibly overcome this limitation and target more covert aspects of gender discrimination. Future research is needed to validate this approach.

Previous studies have shown that different versions of the full-body ownership illusion have various cognitive, emotional, and behavioral consequences. For example, attitudes toward other people change after illusory ownership of their bodies^20^, emotional feelings of social fear^34^ and body dissatisfaction^23,24^ can be modulated by the full-body ownership illusion, and the encoding of episodic memories depends on the embodied first-person perspective^47^. Even the recognition of one’s own face^48–50^, the style of one’s own behavior^51^, and implicit associations with attributes that one used to possess^30^ are flexibly adjusted to the ongoing perception of one’s own body. With regard to gender, it has been shown that the induction of the body-sex-change illusion is possible^15,35^ and that female participants who looked at male avatars from a first-person perspective improved their working memory performance during a stereotype-threatening situation^52^; however, the latter finding is difficult to interpret, as there was no conclusive evidence that the participants felt ownership of the avatar’s body. Our study extends the above literature in three ways: first, by showing that even the supposedly most stable aspects of the psychological sense of self, that is, gender identity, are dynamically updated by the ongoing perception of one’s own body; second, by demonstrating that this updating affects both implicit and explicit beliefs about the self; and third, by clarifying that the illusory ownership of another person’s body not only modifies attitudes toward this person or toward a social group to which this person belongs but also modifies beliefs about the self.

With regard to cognitive mechanisms of the body-related flexibility of self-concept, there are several possible explanations. Embodied cognition theories propose that all concepts are grounded in sensorimotor and situated representations; thus, a change in own body representation, for example, during a full-body ownership illusion, affects conceptual knowledge about the self^20,53^. In turn, predictive processing theories suggest that if the low-level perceptual representation of one’s own body creates a conflict further up in the processing hierarchy, then this conflict is resolved by updating higher-order beliefs about oneself^19,20^. Alternative interpretations focus on cognitive dissonance, self-perception, bodily resonance, or priming^52,54,55^. What the present results add to this complex discussion is the demonstration that the body-sex-change illusion *weakened* implicit associations between the self and the preferred gender category rather than *strengthened* associations between the self and the opposite gender (Fig. 2f), which suggests that the illusion was related to increased cognitive conflict within the existing structure of knowledge about the self that arose as a consequence of the perceived bodily sex change.

We speculate that at the neural level, the fundamental interplay between the perception of one’s own body and gender identity is implemented by functional interactions between the multisensory frontoparietal areas that represent the bodily self^21,22,28^, on the one hand, and medial prefrontal regions that are involved in self-concept representation^56,57^, affective body representations in the insula and anterior cingulate cortex^24^, and higher-order visual representation of the body in the lateral occipital cortex^58,59^, on the other hand. Multisensory representations in the posterior parietal cortex may be particularly important in this respect, as this region is sensitive to the perceived size and shape of one’s own body^24,60^, including waist size^60^, which is likely to be important for the identification of the body’s sex based on secondary sex characteristics. Notably, the pattern of resting-state connectivity in the posterior parietal cortex is altered in transgender individuals compared to cisgender age-matched controls^11^, and a recent study reported that individuals with gender dysphoria display greater cortical thickness in the anterior cingulate cortex and lateral occipital cortex than controls^13^. Interestingly, the lateral occipital cortex, which includes the extrastriate body area – a higher-order visual area that is involved in the processing of images of human body parts^61^ – shows increased activation during body ownership illusions in healthy participants^24,29,58,59^. Future neuroimaging studies could use the present sex-body-swap illusion to perturb the sense of gender identity in cisgender and transgender individuals and investigate how the patterns of activity and functional connectivity within the above fronto-parieto-occipital networks change accordingly.

It is noteworthy that our findings were mainly related to balancing identification with both genders rather than to a “full switch” to the opposite gender. This could be because the perception of one’s own body is not potent enough to completely override the existing sense of own gender or because the body-sex-change illusion in the present study was not induced for long enough. Future studies are needed to reveal the extent to which gender identity could change as a result of a modified body representation and the persistence of these changes over time. Another methodological aspect that is noteworthy is that the body-induced fluidity of gender identity showed relatively large interindividual differences. This variability was related mainly to how vividly the participants experienced the body-sex-swap illusion, which of course makes sense because if there is no change in the representation of one’s body, then there is no reason to update one’s gender identity. Individual differences in the strength of body ownership illusions are generally considered to relate to how brains integrate visual, tactile, and proprioceptive signals^21,62^ depending on the relative weights assigned to different sensory channels and prior knowledge that varies across subjects^25,63^. For example, if more weight is given to vision than to proprioception, the illusion should be stronger, and vice versa. Note that when the illusion is quantified, as in the present study, that is, as the difference in illusion-related ratings between a synchronous illusion condition and an asynchronous control condition, individual differences are unlikely to be related to suggestibility^64^. Moreover, based on our data, we can conclude that variability in the illusion strength was not significantly related to the participants’ sex, age, or baseline feelings of masculinity/femininity (Supplementary Fig. S2 and S3). Finally, it is worth mentioning that our within-subject experimental design allowed us to demonstrate a particularly strong case of gender identity flexibility that occurred for the *same* participants across different body perception contexts.

In sum, the present study shows that there exists a dynamic and automatic link between the perception of one’s own body and different aspects of the sense of own gender. This main finding has important bearings on neurocognitive models of gender identity, as well as on clinical psychology and psychiatry, as the body-sex-change illusion that we report here allows a manipulation of gender identification in nontransgender participants, which offers an unprecedented opportunity to investigate the sense of own gender in a controlled experimental setting. Moreover, people with gender dysphoria who consider surgical and hormonal treatment to adjust their physical appearance to their gender identity could perhaps benefit from future iterations of the body-sex-change illusion, which, combined with virtual reality and 3D body scanners, could allow these patients to experience their “new body” to some extent, before undergoing more permanent procedures.

## Materials and Methods

All participants provided written informed consent before the start of each experiment. The Regional Ethics Review Board of Stockholm approved the studies, and all methods were performed in accordance with the approved guidelines. No participant reported a history of psychiatric illness or neurological disorder, and all had normal or corrected-to-normal vision and hearing. Sample sizes were based on similar previous studies (see Introduction) and our counterbalancing scheme. Data collection was finalized when the planned number of participants was reached. At the end of each experiment, the participants were debriefed and received compensation. All measures used in the study are reported in the manuscript.

## Experiment I

### Participants

Thirty-three naïve healthy adults participated (age: 25±4; 4 left-handed; 15 females). Data from one participant were excluded due to a procedural error (same condition repeated twice).

### Procedure

The participants first rated how masculine or feminine they felt before experiencing any body perception manipulation (baseline; Fig. 1d). The main experiment consisted of four conditions (“synchronous opposite sex”, syncO; “synchronous same sex”, syncS; “asynchronous opposite sex”, asyncO; and “asynchronous same sex”, asyncS), each lasting 3.5 min. During each condition, the participants lay on a bed with their head tilted forward (∼45°) and wore a head-mounted display (HMD; Oculus Rift Development Kit 2, Oculus VR, Menlo Park, CA, USA) so that they could not see their actual body. In the HMD, the participants watched prerecorded 3D videos of a stranger’s body, male or female, that was shown from a first-person perspective. The stranger’s body was continuously stroked on the thighs and abdomen, and the experimenter delivered synchronous (syncO, syncS) or asynchronous (1 s delayed; asyncO, asyncS) touches on the corresponding parts of the participant’s body (Fig. 1a). During each condition, there were three “knife threats” that occurred 1, 2, and 3 min after the beginning of each video (Fig. 1c and 1e). After each condition, the participants took off the HMD, filled out the illusion questionnaire (Fig. 1b), and rated how masculine or feminine they felt during the previous session (Fig. 1d). The order of conditions was counterbalanced across the participants, and the whole experiment lasted ∼30 min (Fig. 1e).

### Prerecorded videos

During filming, a male and a female lay still on a bed, and the experimenter used a 90-cm-long stick with a white plastic ball (diameter 10 cm) attached to its end to deliver strokes to each model’s abdomen, left thigh, or right thigh. The duration of each stroke was 1 s, and each stroke covered ∼20 cm of the model’s body. The time between the end of one touch and the onset of the next touch ranged between 3 and 5 s. The frequency of the strokes was 12 times per minute, and the order of the strokes was pseudorandom (no more than two successive strokes of the same body part). Altogether, 36 strokes (12 to each body part) were delivered during each video. The videos were recorded with two identical cameras (GoPro HERO4 Silver, GoPro, Inc., San Mateo, CA, USA) placed parallel to each other (8 cm apart) just above the models’ heads. The recordings from both cameras were combined in a single frame using Final Cut Pro software (version 7, Apple Inc., Cupertino, CA, USA). Two versions of high-quality 3D videos were created: one for the male and one for the female body. Audio cues were then added to each video. These cues were either congruent with the touches applied in the videos (same body parts, same onset, same duration) or delayed by 1 s. The experimenter listened to these cues during the experiment and applied the touches accordingly. All other aspects were identical in the synchronous and asynchronous videos.

### Knife threats

For each video, we recorded knife-threat events. During these events, a hand holding a knife entered the field of view from above and performed a stabbing movement toward the model’s body (Fig. 1c). The knife stopped just before hitting the body, changed direction (−180°), and exited the field of view in the same way that it had entered. The whole event lasted 2 s. After each knife threat, there was a 10 s pause in the video with no strokes delivered.

### Visuotactile stimulation during the experiment

The experimenter listened to audio cues from the videos (see earlier) and accordingly applied touches to the participant’s body. These cues were played via headphones, so the participants could not hear them. The number, order, type, length, velocity, and frequency of the strokes applied during the experiment precisely followed the strokes in the prerecorded videos (see earlier). To deliver the touches, the experimenter used the same white ball attached to a stick that had been used during the video recordings.

### Illusion questionnaires

The subjective experience of the full-body ownership illusion was quantified with a questionnaire that began with an open-ended sentence (“During the last session, there were times when…”). This sentence was followed by three illusion statements that quantified the explicit feeling of body ownership (I1; Fig. 1b) and the sensation of touch directly on the stranger’s body (I2 and I3; Fig. 1b), which are considered two core elements of the multisensory full-body illusion^16,17^. Apart from the illusion statements, the questionnaire included four control statements (C1-C4; Fig. 1b) that were added to control for potential task compliance or suggestibility effects. In the questionnaire that was handed to the participants, the items were listed in the following order: C1, I1, C2, I2, C3, C4, I3. The participants marked their responses on a scale from −3 (“strongly disagree”) to +3 (“strongly agree”).

### Skin conductance responses

The skin conductance response reflects increased sweating attributable to the activation of the autonomic nervous system^65^. When one’s body is physically threatened, the threat triggers emotional feelings of fear and pain anticipation that are associated with autonomic arousal. This arousal can be registered as a brief increase in skin conductance a few seconds after the threat event. Increased threat-evoked skin conductance responses during an illusion condition compared to a well-matched control condition are often used as an index of body ownership in body illusion paradigms^15,21,29^. In the current experiments, data were recorded continuously with Biopac system MP150 (Biopac Systems Inc., Goleta, CA, USA) and AcqKnowledge software (version 3.9). The following parameters were used: sampling rate=100 Hz, low-pass filter=1 Hz, high-pass filter=DC, gain=5 μS/V, and CAL2 scale value=5. Two Ag-AgCl electrodes (model TSD203, Biopac Systems Inc., Goleta, CA, USA) were placed on the volar surfaces of the distal phalanges of the participants’ left index and middle fingers. Isotonic paste (GEL101; Biopac Systems Inc., Goleta, CA, USA) was used to improve the skin contact and recording quality. At the beginning of the experiment, we asked the participants to take the deepest breath possible and hold it for a couple of seconds. In this way, we tested our equipment and established a near maximum response for each participant. The timings of the threat events were marked in the recording file by the experimenter by pressing a laptop key immediately after the threat occurred.

## Experiment II

### Participants

Sixty-four naïve volunteers participated (age: 27±5; all right-handed; 32 females).

### Procedure

The participants first completed a practice IAT (20 trials). The main study consisted of the same four conditions as those used in Experiment I. After the initial phase of just watching the videos and feeling touches (30 s), the participants started the first IAT block (Fig. 2b and 2c). The IAT stimuli were presented via headphones (Spectrum, Maxell Europe Ltd., Berkshire, UK), and the participants used a wireless computer mouse, held in the right hand, to indicate their responses. During each condition, the participants saw two “knife threats”, one in the middle and one at the end of each condition (Fig. 2c). After each condition, the participants filled out the same illusion questionnaire used in Experiment I (Fig. 1b). The order of the conditions was counterbalanced, and the whole study lasted ∼1 h (Fig. 2c).

### Prerecorded videos

The videos were prepared analogously to those in Experiment I, but a different male and female were filmed to assure that our results were not driven by a certain body type or clothing style of the models (Fig. 2a). Strokes were applied to three body parts: abdomen, left thigh, and right thigh. The abdomen strokes were either single or double (1 s apart). The duration of each stroke was 1 s, and each stroke covered ∼20 cm of the model’s body. The time between the offset of one touch and the onset of the next touch ranged from 3 to 6 s. The frequency of the strokes was 12 times per minute. Different touches were delivered in a pseudorandom sequence, with no more than three successive strokes of the same body part. Altogether, 88 touches (22 to each body part) were applied in each video. The videos were recorded with Infinity cameras (1080p Full HD, CamOneTec, Delbrück, Germany) and prepared in the same way as in Experiment I. In the synchronous videos, audio cues were matched with the touches applied in the videos, whereas in the asynchronous videos, the cues were delayed by 1 s and prompted different body parts. Altogether, we created four versions of high-quality 3D videos, each lasting 7 min 5 s.

### IAT

We used the auditory version of the brief gender identity IAT^40,41^. The instruction for one block was as follows: “The test will start in a few seconds. Please listen to the instructions. Try to go as fast as possible while making as few mistakes as possible. If the word belongs to the categories *female* or *self*, press left. If the word does not belong to these categories, press right. The test will begin now.” The instruction for the other block differed only with regard to category assignment: “If the word belongs to the categories *male* or *self*, press left. If the word does not belong to these categories, press right.” The key assignment remained the same for a given participant but was counterbalanced between the participants. The order of the IAT blocks was counterbalanced in the same way. The stimulus set consisted of twenty words (Fig. 2b) that were read by an English native speaker. The volume of each word sound was adjusted using Audacity software (the “normalize” effect; version 2.1.2, http://www.audacityteam.org). Each word was edited to have a duration similar to that of other words. Please note that physical differences between the stimuli cannot explain the main IAT effect (different reaction times in the congruent and incongruent blocks), as both blocks used exactly the same stimuli, just with different instructions. The participants had a maximum of 3 s to provide a response (time from the stimulus onset to the end of each trial). If no key was pressed within this time or the wrong key was pressed, the participants heard a “wrong” feedback sound. Each IAT block consisted of 60 trials (three repetitions of all 20 words) presented in random order. The IAT procedure was self-paced; that is, the next trial started as soon as the participant responded in the previous trial (maximum duration of one block ∼3 min). Presentation software (Neurobehavioral Systems Inc., Albany, CA, USA) was used to present the stimuli and record responses.

### Knife threats

These events were recorded in the same way as in Experiment I, but this time, there were only two threats per condition (Fig. 2c; one after the first IAT block, in the middle of each video, and one after the second IAT block, at the end of each video). Because the IAT was initiated at the same time as the prerecorded videos, we could use triggers from Presentation software to automatically flag the onsets of the knife threats in the skin conductance recording files.

## Experiment III

### Participants

Forty-five naïve healthy volunteers participated (age: 26±5; all right-handed; 22 females). One participant was excluded because he did not complete one of the questionnaires.

### Procedure

The study lasted ∼35 min and comprised two conditions: syncO and asyncO (Fig. 3a and 3b). Each condition lasted 14 min 10 s. After each condition, the participants filled out the illusion questionnaire (the same as in Experiment I) and the Bem-Sex-Role-Inventory; BSRI^42,43^ (see further). The order of the conditions was counterbalanced across participants (Fig. 3b).

### Prerecorded videos

The videos were prepared analogously to those in Experiments I and II. Four types of strokes (single abdomen, double abdomen, left thigh, and right thigh) were applied. The duration of each stroke was 1 s, and each stroke covered ∼20 cm of the model’s body. The time between the offset of one touch and the onset of the next touch ranged from 2 s to 10 s. The frequency of the strokes was 12 times per minute. Different touches were delivered in a pseudorandom sequence, with no more than three successive strokes of the same body part. Altogether, 160 touches (40 to each body part) were applied in each video. Infinity cameras (1080p Full HD, CamOneTec, Delbrück, Germany) were used to record the videos. Audio cues were matched to touches in the synchronous videos and delayed by 1 s in the asynchronous videos.

### BSRI

After each condition, the participants filled out a version of the BSRI^42,43^. This questionnaire contained 5 stereotypically masculine and 5 stereotypically feminine personality traits (Fig. 3c). Using a 7-point Likert scale (1 – “not at all”; 7 – “very much”), the participants rated how well each trait described them. Half of the traits were rated after the first condition and the other half after the second condition. The order of the BSRI versions was counterbalanced.

## Analyses

### Analysis of illusion questionnaires

In all experiments, for each participant, we calculated “illusion scores” as differences between the average illusion (I1-I3) and control (C1-C4) ratings in each condition (Fig. 1b). The illusion scores were analyzed with the linear mixed model: score ∼ 1 + body + synchrony + body*synchrony + 1|id. The fixed effects of “body” and “synchrony” had two levels (same sex vs. opposite sex and synchronous vs. asynchronous, respectively). The “1|id” refers to the random intercept, which accounted for the general variability of the illusion scores between participants. The ratings of individual statements are shown in Supplementary Fig. S5. The fixed effect of “ownership” used in some of the analyses (Fig. 1 h, Fig. 2f, and Fig. 3f) was operationalized as the difference between I1 ownership ratings in syncO – asyncO. The participants who experienced the body-sex-change illusion most vividly were selected with the median-split method applied to I1 ownership ratings: syncO – asyncO. The analyses performed on this subgroup of participants (Fig. 1i and Fig. 2g) were performed mainly for display purposes and to complement the main analyses using continuous scores.

### Analysis of skin conductance responses

Each response was measured as a difference between the maximum and minimum values during the 0 to 6 s period after each knife threat. Responses below 0.02 μS were treated as zeroes but were included in the analysis of magnitude skin conductance responses^65^. Statistical outliers were identified with the ±1.5 interquartile criterion and removed from the dataset (16% and 6% of values in Experiments I and II, respectively). Keeping the outliers did not change the overall pattern of the skin conductance results (main effect of synchrony; Experiment I: *b*=0.13; *SE*=0.03; *t*=3.79; *P*<0.005; N=32; Experiment II: *b*=0.21; *SE*=0.06; *t*=3.54; *P*<0.005; N=64; linear mixed models; two-sided). The data were analyzed with the following linear mixed model: response ∼ 1 + body + synchrony + body*synchrony + repetition + 1|id (see earlier for explanation of factors). The fixed effect of “repetition” included values from 1 to 12 (Experiment I) or from 1 to 8 (Experiment II) and indicated how many knife threats had already occurred in the study. It is well established that skin conductance responses decrease with subsequent threats^65^, and we found this habituation effect as well (main effect of repetition; Experiment I: *b*=0.29; *SE*=0.03; *t*=8.7; *P*<0.005; Experiment II: *b*=0.59; *SE*=0.05; *t*=12.63; *P*<0.005; linear mixed models; two-sided; N=32 and N=64, respectively). In fact, the responses decreased exponentially with subsequent knife threats, so a transformed repetition number (1/n) substantially improved the fit of the linear model to the data (Experiment I: χ^21^=4.36; *P*<0.005; Experiment II: χ^21^=37.26; *P*<0.005; linear mixed models; two-sided; N=32 and N=64, respectively).

### Analysis of masculinity/femininity ratings

The data were analyzed with the following linear mixed model: rating ∼ 1 + body + synchrony + body*synchrony + 1|id (see earlier). Follow-up pairwise comparisons were performed with the following model: rating ∼ 1 + condition + 1|id. For the analysis presented in Fig. 1h, we corrected all values in the following way: rating from a given condition = rating from this condition – average of ratings from all conditions for a given participant (we did not divide by the standard deviation because it was zero for some participants). This correction accounted for overall between-subject differences in feelings of masculinity or femininity. The analysis shown in Fig. 1h used the following model: rating ∼ 1 + synchrony*body*ownership + 1|id (all main effects and two-way interactions were also included). Follow-up analyses for each condition were conducted with linear regression models: rating ∼ 1 + ownership + 1|id.

### Analysis of IAT

The data included only correct trials in which reaction times were longer than 200 ms and shorter than 1500 ms (95.5% of all trials; n=29,147). Reaction times were analyzed with the following linear mixed model: reaction time ∼ 1 + body*synchrony*block + 1|id + 1|item (all main effects and two-way interactions were also included). The fixed effect of “block” had two levels: incongruent and congruent. The random-intercept “1|item” referred to the word that was presented in a given trial (“self, “they”, “Linda”, etc.). Follow-up comparisons for each condition used the following linear mixed model: reaction time ∼ 1 + block + 1|id + 1|item. For the analysis presented in Fig. 2f, we used the following model: reaction time ∼ 1 + body*synchrony*block*ownership + 1|id + 1|item. The follow-up analyses for each condition used the following model: reaction time ∼ 1 + block*ownership + 1|id + 1|item. The following model was used to analyze data from the congruent and incongruent blocks in the syncO condition: reaction time ∼ 1 + ownership + 1|item (Fig. 2f).

### Analysis of BSRI

For each participant in each condition, we calculated the stereotype-congruent (c) and stereotype-incongruent (i) scores as average ratings of masculine and feminine traits. To account for overall between-subject differences, we standardized each score in the following way: a given score = (this score – average of all scores for a given participant) / standard deviation for a given participant. These scores were compared within the syncO and asyncO conditions using the following linear mixed model: score ∼ 1 + stereotype-congruency + 1|id (Fig. 3e). The relationship between stereotypical bias reduction (BSRI scores: asyncO^c-i^ – syncO^c-i^) and illusory ownership of the opposite-sex body (I1 ownership ratings: syncO – asyncO) was tested with the following linear regression model: bias reduction ∼ 1 + ownership.

### Meta-analysis

For each participant in each experiment, the degree of gender identity updating was calculated as the difference between gender preference in the syncO condition versus the control conditions. Specifically, in Experiment I, this updating score corresponded to the following difference between masculinity/femininity ratings: avg. (syncS, asyncS, asyncO) – syncO. In Experiment II, this updating was calculated as the difference between reaction times: avg. (syncS^i-c^, asyncS^i-c^, asyncO^i-c^) – syncO^i-c^, where “i-c” is the difference in milliseconds between the incongruent and congruent IAT blocks. Finally, in Experiment III, this updating score was calculated as the difference between BSRI ratings: asyncO – syncO. To compare these scores across experiments (different scales), we standardized each score within a given experiment; we subtracted a group mean and divided the result by the standard deviation from a particular experiment.

### General statistical information

All statistical analyses were performed in RStudio and R software (version 3.3.3, The R Foundation for Statistical Computing, https://www.r-project.org). Linear mixed models were estimated using the “lme4” package. *P*-values were obtained with the bootstrapping method by comparing a given parameter value to its null distribution derived from iterative resampling of the original dataset (“boot” package; 10000 simulations).

## Acknowledgments

This study was funded by the Swedish Research Council, Torsten Söderbergs Stiftelse, Göran Gustafsons Stiftelse, StratNeuro, and the European Commission (MSCA fellowship awarded to P.T.). We want to thank all the participants and Martti Mercurio for important technical support.

## Author Contributions

P. T. and H. H. E. designed the study and wrote the manuscript. P. T. and J. F. collected and analyzed the data. All authors provided revisions and approved the final version of the manuscript for submission.

## Data availability

Group-level data related to the analyses presented in figures from this study are included as Supplementary Information files. We do not have ethics approval to make the raw data from individual subjects publicly available.

