## Supplementary Information for "Fluidity of gender identity induced by illusory body-sex change"

P. Tacikowski, J. Fust, and H. H. Ehrsson

**This document includes the following:**

Supplementary Figures S1-S5

Supplementary Datasets Legend

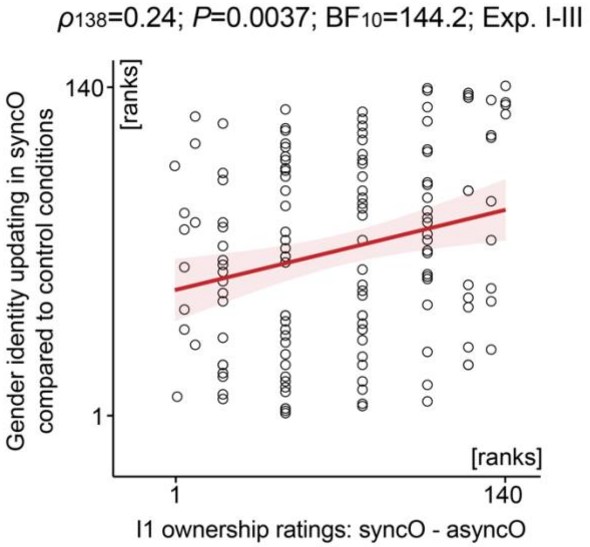


**Fig. S1.** Meta-analysis on data from all experiments confirmed that illusory ownership of the opposite-sex body was associated with increased updating of the sense of own gender (Spearman's correlation; two-sided; N=140; $BF_{10}$ indicates a Bayes factor in support of the alternative hypothesis).

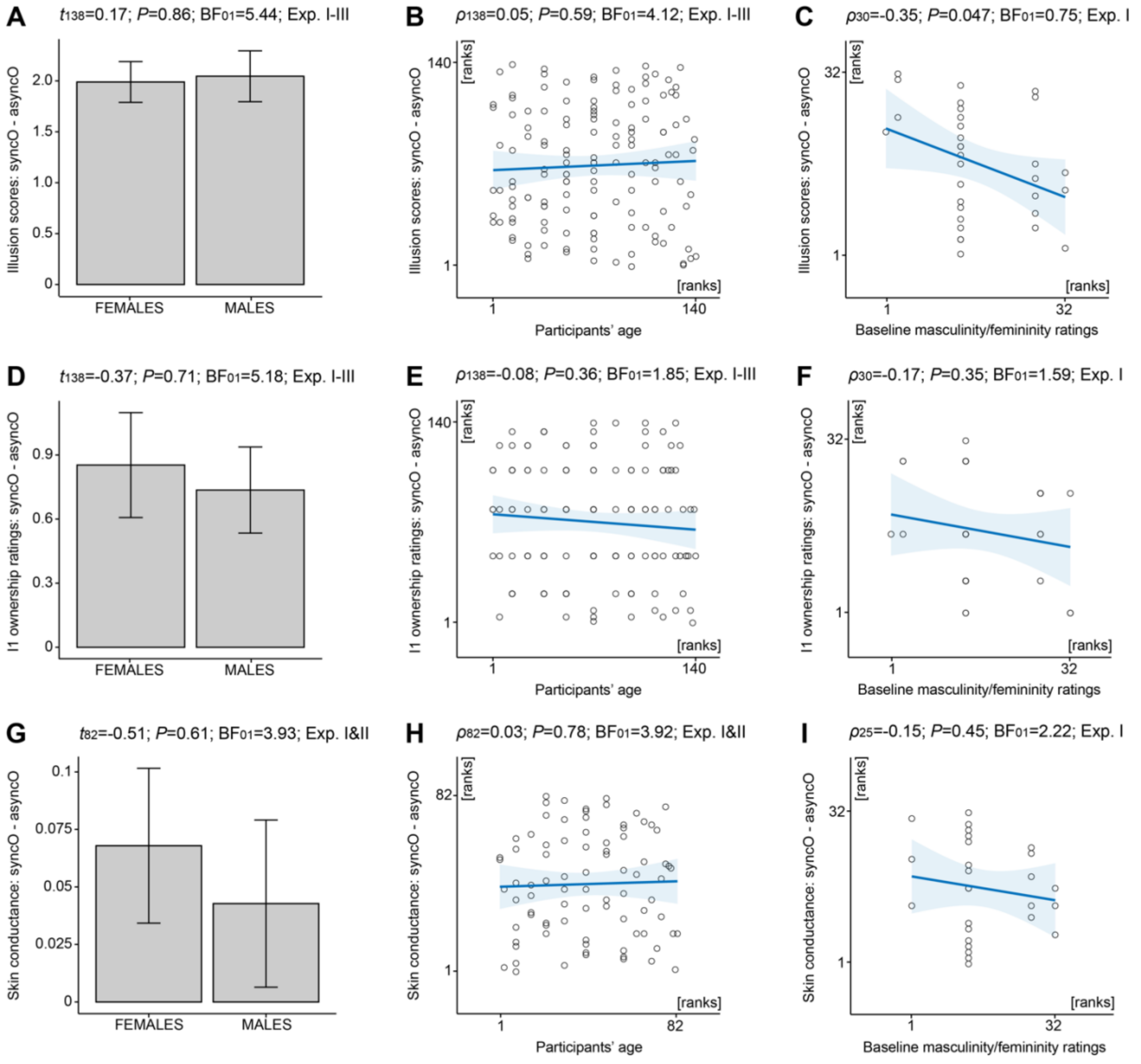


**Fig. S2. The strength of the body-sex-change illusion in syncO was similar among male and female participants, and there was no consistent significant modulation of the illusion strength by the participants' age or baseline masculinity/femininity ratings.** The strength of the full-body illusion was measured as the syncO – asyncO difference between illusion scores **(A-C)**, I1 ownership ratings **(D-F)**, and skin conductance responses **(G-I)**. In panels C, F and I, the higher the rank on the x-axis, the higher the rating of feeling masculine (biological males) or feminine (biological females) during baseline (Fig. 1d). Continuous variables were analyzed with Spearman's correlation tests. The effect of participants' sex was tested with the independent-samples t-tests. $BF_{01}$ indicates Bayes factors in support of the null hypotheses. All *P*-values are two-sided. Bar plots show means±*SE*.

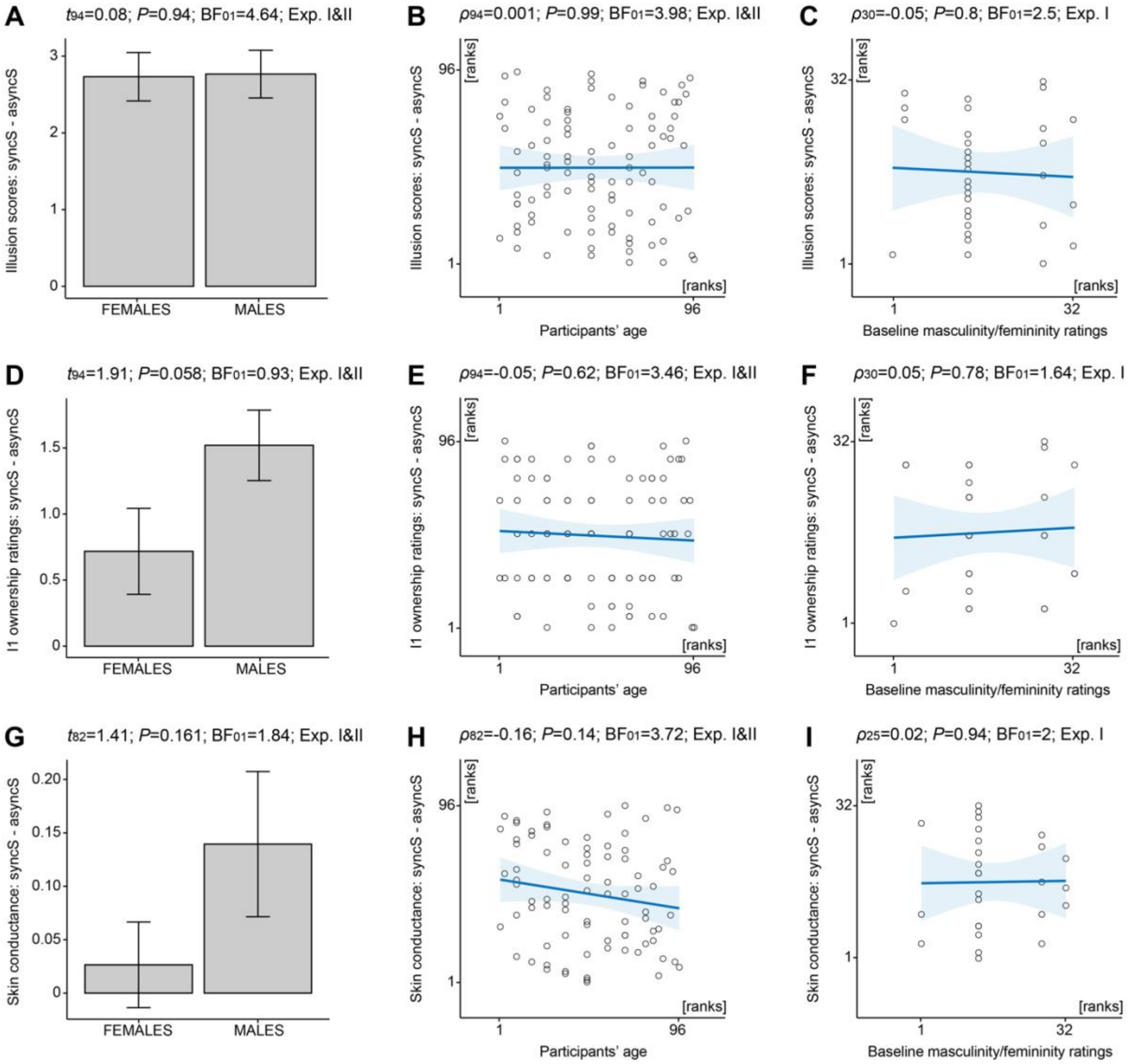


**Fig. S3. The strength of illusory ownership of the same-sex stranger's body (syncS) did not differ significantly between males and females, and there was no significant relationship between the illusion in syncS and the participants' age or baseline masculinity/femininity ratings.** The strength of the full-body illusion in syncS was measured as the syncS – asyncS difference between illusion scores **(A-C)**, I1 ownership ratings **(D-F)**, and skin conductance responses **(G-I)**. Continuous variables were analyzed with Spearman's correlation tests. The effect of participants' sex was tested with the independent-samples t-tests. $BF_{01}$ indicates Bayes factors in support of the null hypotheses. All *P*-values are two-sided. Bar plots show means±*SE*.

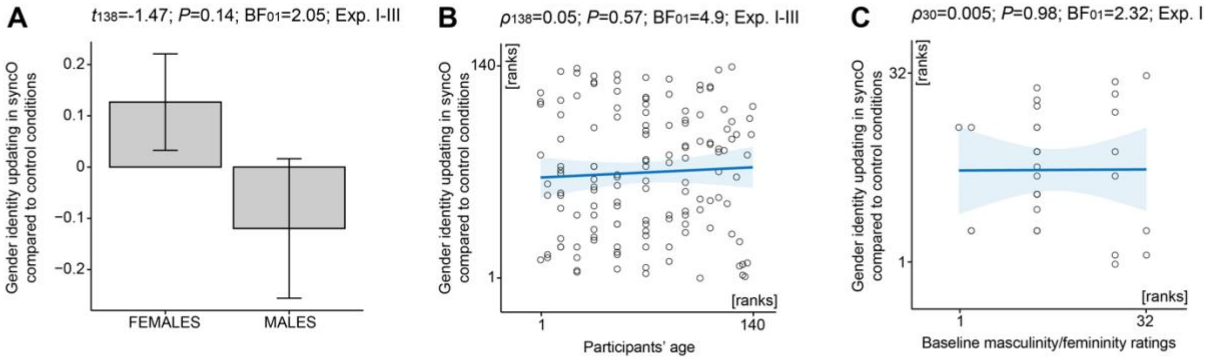


**Fig. S4.** The degree of gender identity updating was not significantly different across males and females (A), and there was no significant relationship between the degree of this updating and the participants' age (B) or the participants' baseline masculinity/femininity ratings (C). Age and masculinity/femininity ratings were analyzed with Spearman's correlation tests. The effect of participants' sex was tested with the independent-samples t-test. $BF_{01}$ indicates Bayes factors in support of the null hypothesis. All *P*-values are two-sided. Bar plots show means±*SE.*

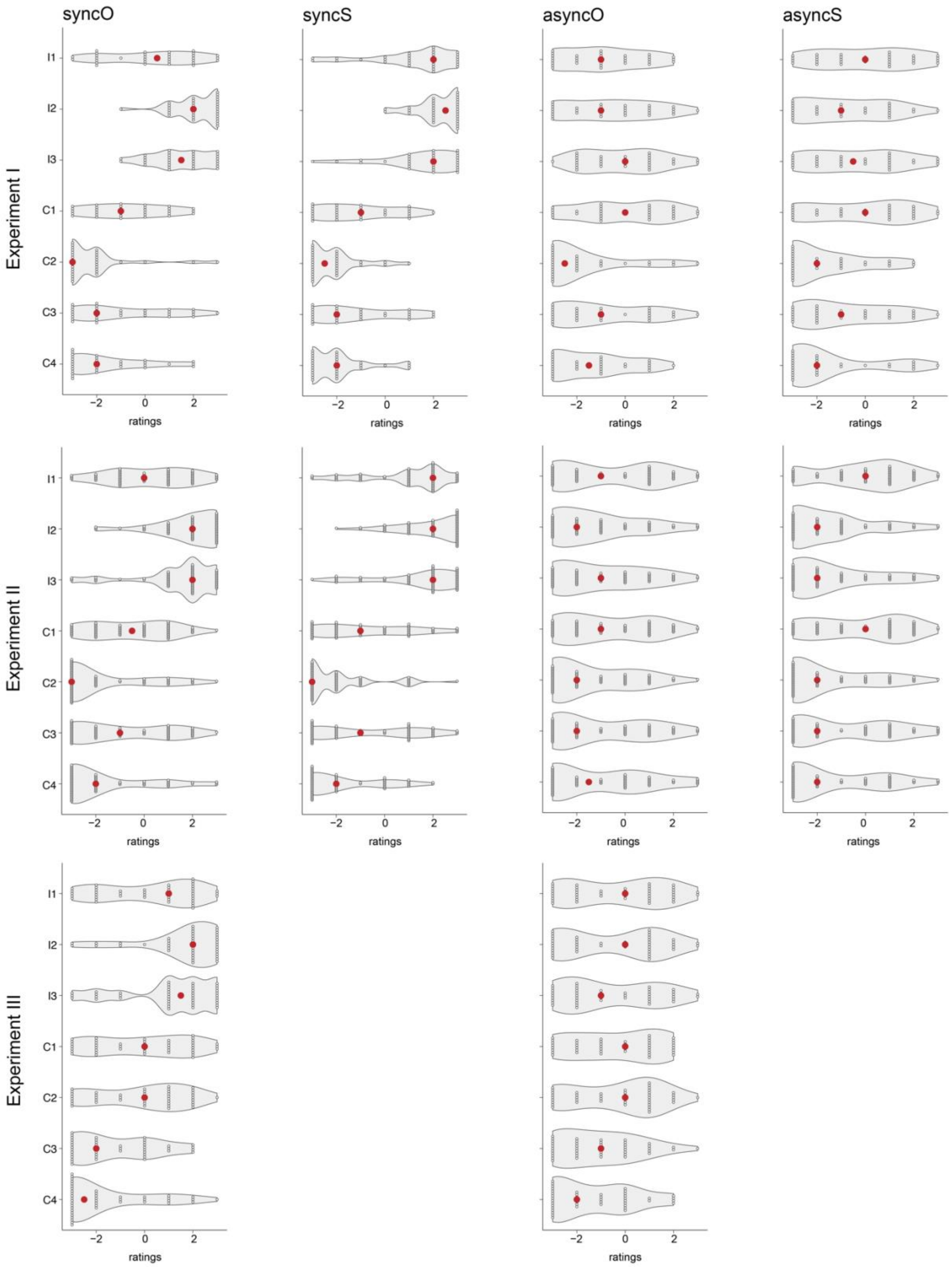


**Fig. S2. Illusion questionnaire results.** Red dots represent medians, small white dots are individual ratings, and "clouds" are probability density of ratings at different values. I1:I3 are illusion items, and C1:C4 are control items.

**Datasets legend:**

**Dataset_1:** Experiment I: Condition order, illusion questionnaire ratings, and masculinity/femininity ratings.

**Dataset_2:** Experiment II: Condition order, block order, and illusion questionnaire ratings.

**Dataset_3:** Experiment III: Bem-Sex-Role-Inventory (BSRI) ratings and illusion questionnaire ratings.
